# *Drosophila* Cocaine Avoidance is Mediated by Peripheral Bitter Gustatory Neurons

**DOI:** 10.1101/2022.06.22.497211

**Authors:** Travis J. Philyaw, Iris Titos, Pearl N. Cummins, Aylin R. Rodan, Adrian Rothenfluh

## Abstract

Genetic variation contributes to heterogeneity in the prevalence of complex disorders such as addiction. The genetic risk for developing a substance use disorder can vary between drugs. The estimated heritability rate of cocaine addiction is 72%, higher than any other drug. Despite recognition of this significant genetic component, little is known about the specific genes and mechanisms that lead to the development of cocaine addiction. *Drosophila* is an effective model organism for identifying the genes that underlie complex behaviors, including addiction. While *Drosophila* exposed to cocaine display features of intoxication similar to those observed in mammals, there is currently no model of cocaine self-administration in flies. Because cocaine is a natural insecticide, we wondered if *Drosophila* might naively avoid it through bitter chemosensory detection. To answer this question, we performed cocaine consumption and preference assays comparing wild-type flies and bitter-taste mutants. Our results demonstrate that *Drosophila* detect and avoid cocaine through bitter sensing gustatory neurons, and that this process requires gustatory receptor 66a (Gr66a). Additionally, we identify a peripheral mechanism of avoidance through cocaine detection with *Drosophila* legs. Our findings reveal that preingestive mechanisms of toxin detection play a significant role in *Drosophila* cocaine avoidance and provide evidence that disrupting gustatory perception of cocaine is essential for self-administration and, therefore, developing a model of self-administration in *Drosophila*.

## Introduction

Cocaine is a psychostimulant that elevates dopamine and mood in humans (Volkow et al., 2009). The reinforcing effects of cocaine intoxication contribute to its addictive potential, and national and international surveys on drug use suggest cocaine is used by more than 20 million individuals globally and estimate that more than one million individuals in the United States struggle with cocaine use disorder (SAMHSA, 2019; UNODC, 2021). Family studies suggest that the risk for developing drug addiction has a heritability component of around 50% (Kendler et al., 2006), with cocaine use disorder having the highest heritability estimate at around 72% (Goldman et al., 2005). Despite this information, few studies have been able to identify novel genes and proteins involved in the development of cocaine addiction. Due to the heterogeneity in the population and its multigenic basis, a large sample size is required to identify these genes, which is challenging in human studies (Ducci and Goldman, 2012). In contrast, model organisms such as *Drosophila melanogaster* provide a way to study the genes that underlie complex behaviors, including addiction, in homogenous populations with a virtually unlimited sample size (Narayanan and Rothenfluh, 2016). Moreover, *Drosophila* is a proven translational model with orthologs for most disease-causing genes in humans (Bier, 2005), including addiction (Lathen et al., 2020).

Like mammals, *Drosophila* exposed to cocaine display dose-dependent changes in motor behavior that include increased grooming, locomotion, stereotypy, and erratic movements, as well as akinesia, seizures, and death at higher doses (McClung and Hirsh, 1998). Cocaine also reduces sleep in *Drosophila* (Lebestky et al., 2009), as it does in mammals (Bjorness and Greene, 2021). Therefore, *Drosophila* show face-valid responses to cocaine. Studies have compared the relative consumption of cocaine between *Drosophila* genotypes and measured changes in cocaine consumption over time (Highfill et al., 2019; Baker et al., 2021; Kanno et al., 2021); however, there is currently no *Drosophila* model for preferential self-administration of cocaine.

Cocaine has been identified as an insecticide (Nathanson et al., 1993), and flies naively avoid cocaine. Identifying barriers to cocaine consumption in *Drosophila* will support the development of a fly model of cocaine abuse disorder. A model for cocaine use disorder in flies would enable the identification of genes that underlie cocaine addiction using high-throughput behavioral experiments that are prohibitive in mammals. Here we show that *Drosophila* sense cocaine as an aversive compound and that they display naïve cocaine avoidance in acute assays of consummatory preference. We find that this avoidance is mediated by bitter-sensing gustatory neurons and demonstrate that flies taste cocaine using G-protein coupled receptors, thereby identifying a peripheral mechanism for cocaine detection and avoidance in *Drosophila*.

## Results

### Cocaine is Aversive to *Drosophila*

To determine whether *Drosophila* would consume cocaine, we starved flies for 6 hours, then measured feeding during a 4-minute assay using a 340mM sucrose solution supplemented with different concentrations of cocaine (3mM, 10mM, 15mM). Blue dye was added to the food to track consumption (Figure 1a), and we observed a dose-dependent decrease in the volume consumed per fly that ate (Figure 1b). Flies offered sucrose without cocaine consumed the largest amount of solution, with a median volume of more than 185nL per fly (Figure 1b). In contrast, the median volume ingested per fly that ate decreased more than 2-fold for flies fed 3mM cocaine, more than 12-fold for flies fed 10mM cocaine, and over 35-fold for flies fed 15mM cocaine (Figure 1b). The results of our blue-dye feeding assay suggest that wild-type flies have an innate dose-dependent aversion to cocaine, which reduces consumption when added to sucrose at concentrations of 3mM or higher in this acute feeding assay. The concentrations of cocaine published in other feeding experiments range from 10uM to 10mM (Kanno et al., 2021). The concentrations used in assays with vaporized free-base cocaine are much higher, with a “moderate” dose considered 5-10ul of a solution at a concentration of approximately 50mM (McClung and Hirsh, 1998; Bainton et al., 2005). While comparable to doses of cocaine administered in other publications, we wondered whether the concentrations of cocaine used in our experiments were too high to assay consumption effectively. Notably, at doses over 3mM cocaine, flies in our experiment began to exhibit seizures (data not shown), suggesting reduced sucrose consumption might be partially due to cocaine-related incapacitation.

**Figure 1.**
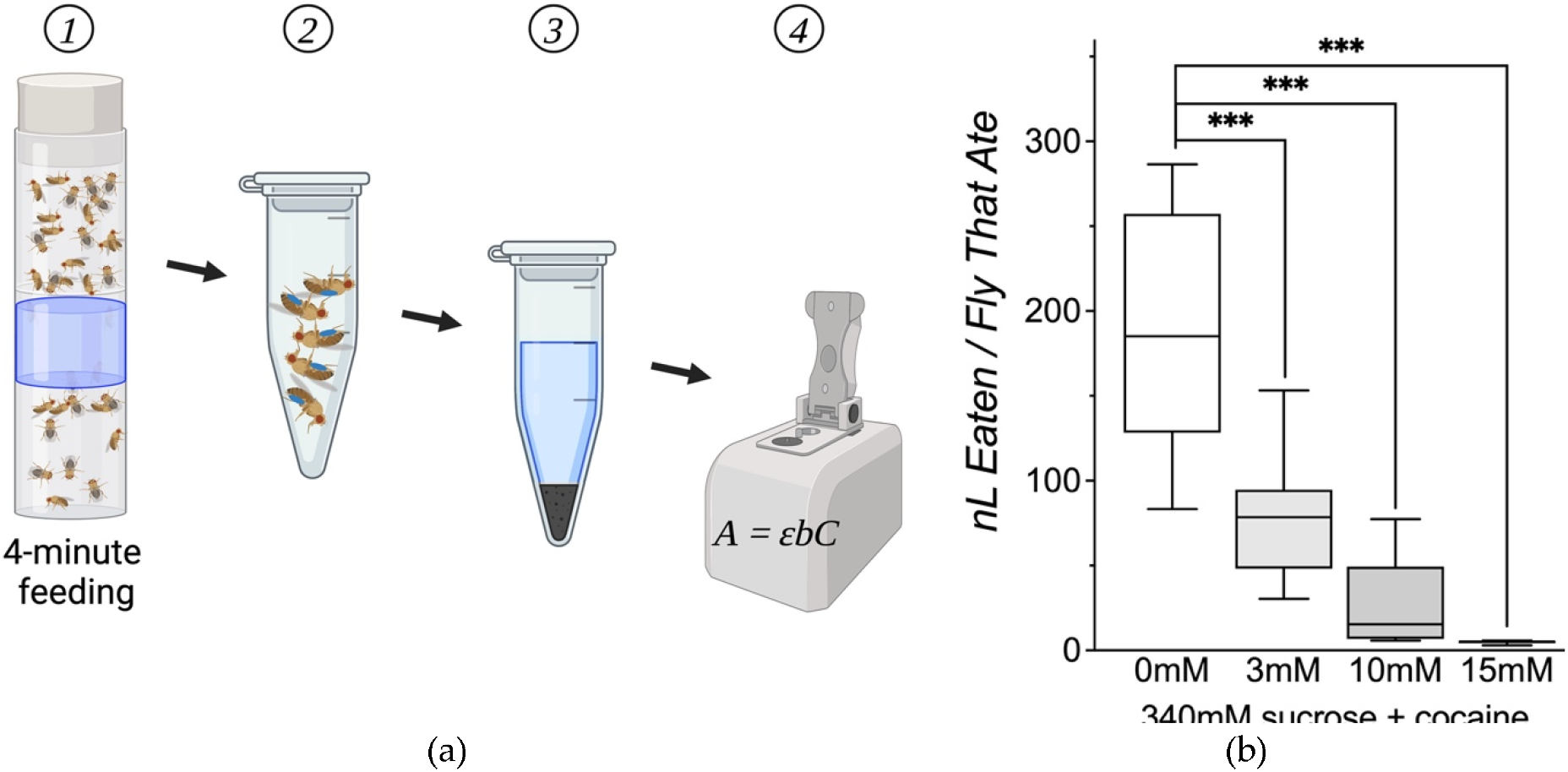
Cocaine Reduces *Drosophila* Sucrose Consumption. **(a)** A schematic of the blue-dye feeding assay of consumption depicting (1) the 4-minute feeding (2) selection of flies that ate → (3) extraction of blue dye → (4) quantification of the volume ingested per fly that ate based on measurement of dye concentration (4). **(b)** The effect of cocaine concentration on median volume consumed (nL) per fly during a 4-minute assay of feeding after a 6-hour wet starvation period. The median consumption per fly that ate is highest for flies offered 340mM sucrose (185nL, n=16 groups of 5 flies). Median consumption per fly that ate is significantly reduced for sucrose solutions containing cocaine, decreasing to 79nL with 3mM cocaine (\*\*\**P* < 0.0004, n = 8 groups of 5 flies, two-tailed Mann-Whitney), 15nL with 10mM cocaine (\*\*\**P* < 0.0001, n = 8 groups of 5 flies, two-tailed Mann-Whitney), and 5nL with 15mM cocaine (\*\*\**P* < 0.0001, n = 6 groups of 5 flies, two-tailed Mann-Whitney). Here, and in subsequent figures, boxplots show the interquartile range, and the line inside each boxplot represents the median. The lower and upper bounds of each box mark the 25th and 75th percentile, respectively. Whiskers indicate the 10th and 90th percentile. Illustrations created with BioRender.com.

To expand on these results, we assayed flies at lower concentrations of cocaine using a fluorometric reading assay of preference (FRAP) (Peru y Colón de Portugal et al., 2014). In contrast to the dye feeding assay, where flies are only offered a single feeding solution, they are given a choice of two solutions for 30 minutes in the FRAP. Each solution in the FRAP is marked with a distinct fluorescent dye, and a fluorescent plate reader is used to measure the relative ingestion of the dyes in whole flies after feeding. These results are used to generate a preference index, representing the relative consumption of each feeding solution (Figure 2a). Preference scores in the FRAP range from −1 to +1, indicating whether flies exhibit avoidance (−) or preference (+) for a tested compound. To determine how *Drosophila* respond to lower concentrations of cocaine, we again starved flies for 6 hours, then offered them a choice between 340mM sucrose and 340mM sucrose supplemented with different concentrations of cocaine (0.3mM, 0.6mM, 1.5mM, or 4.5mM). We observed naïve aversion to cocaine at every concentration tested (Figure 2b). Taken together, the results of our consumption and preference assays show that cocaine is aversive to flies, even at concentrations as low as 0.3mM.

**Figure 2.**
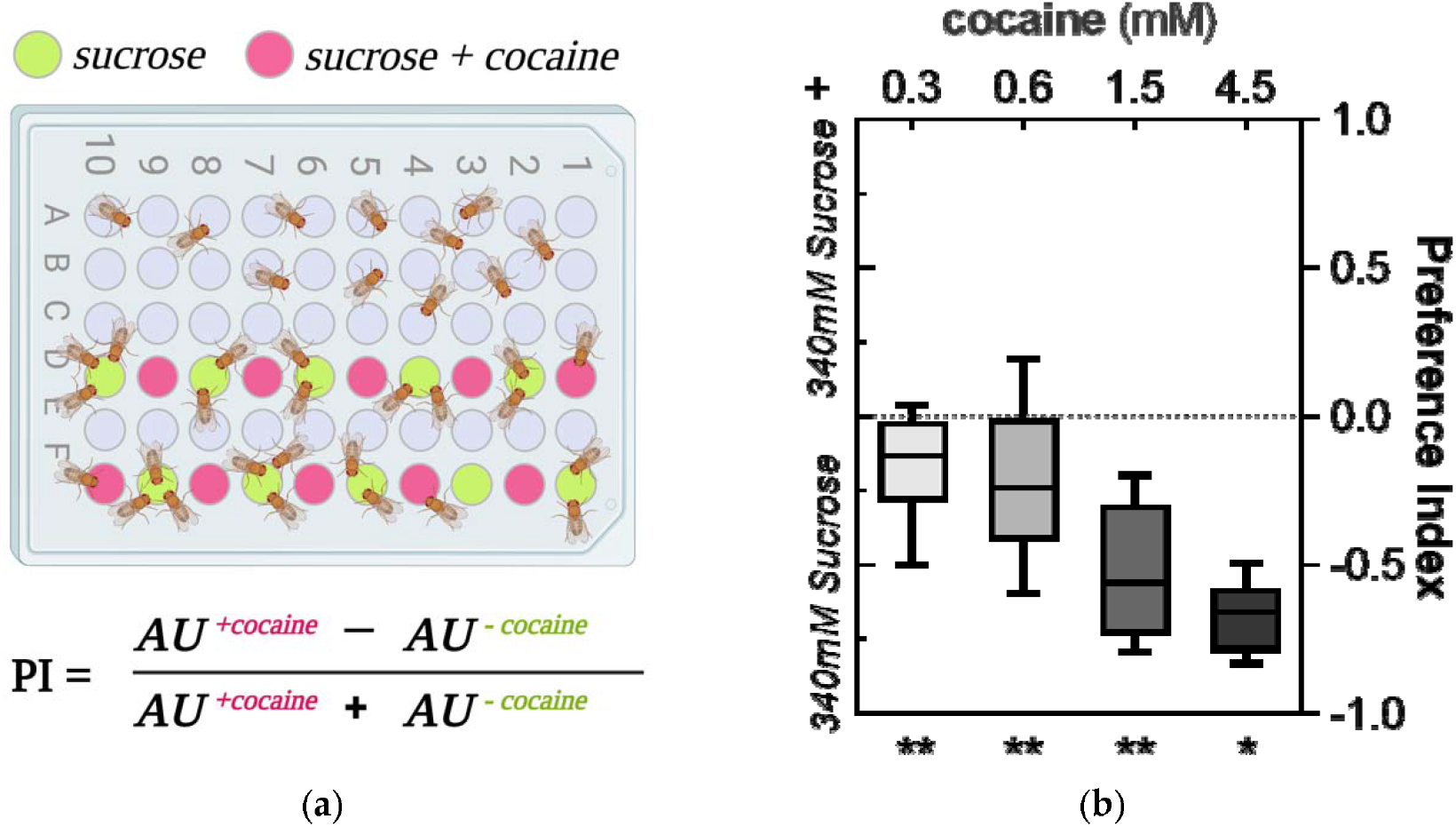
*Drosophila* Avoid Cocaine in a Two-choice Fluorescence Reading Assay of Preference. **(a)** A schematic of preference quantification in one of the two color counter-balanced conditions of the Fluorescense Reading Assay of Preference (FRAP). **(b)** Cocaine is aversive to flies in a 30-minute FRAP. All flies were wet starved for 6 hours then allowed to choose between fluorescently labeled color counter-balanced solutions containing 340mM sucrose or 340mM sucrose supplemented with cocaine. Cocaine produces a dose-dependent decrease in preference, which significantly deviates from a score of 0 (no preference) for sucrose solutions containing 0.3 (*** *P* < 0.001, n = 46 sets of 3 flies), 0.6 (*** *P* < 0.001, n = 47), 1.5 (*** *P* < 0.001, n = 16), and 4.5mM cocaine (* *P* = 0.031, n = 8; one-sample Wilcoxon signed rank test with *P* values Bonferroni-adjusted for multiple testing).

### Cocaine Aversion Involves *Drosophila* Gustatory Neurons

Cocaine is a naturally occurring insecticide found in the leaves of the coca plant and is known to inhibit feeding behavior in other insects (Nathanson et al., 1993). Since most phytotoxins are alkaloids perceived as bitter (Dweck and Carlson, 2020), and flies avoid ingesting bitter substances, we wondered whether bitter chemosensory neurons play a role in *Drosophila* cocaine avoidance. *Drosophila* chemosensory neurons are housed in sensilla found on their sensory organs (Ling et al., 2014). These sensilla contain multiple taste neurons, with each neuron tuned to detect a specific taste modality (Freeman and Dahanukar, 2015). The tuning to detect various tastants is determined by the complement of gustatory receptor (Gr) subunits expressed in different taste neurons (Weiss et al., 2011). While 28 bitter-tasting Grs have been identified, only a subset is expressed in each bitter gustatory neuron (bGRN) (Ling et al., 2014). Some gustatory receptor subunits are narrowly-tuned and respond to a limited number of bitter compounds, while others are widely-tuned and respond to many bitter compounds. A small subset of these widely-tuned, bitter-sensing Gr proteins, referred to as core bitter receptors, are expressed almost ubiquitously in bGRNs (Weiss et al., 2011). One such highly expressed, core bitter receptor protein commonly used to label bGRNs is Gr66a (Weiss et al., 2011).

To determine whether bitter perception contributes to *Drosophila* cocaine avoidance, we abolished bitter taste in flies with the GAL4/UAS expression system (Brand and Perrimon, 1993). We used the Gr66a driver (Dunipace et al., 2001) to promote the expression of an inward rectifying potassium channel (Kir) (Hodge, 2009) that electrically silences neurons, therefore inhibiting bitter sensing in flies. We starved flies for 6 hours and then used the FRAP to determine whether bitter perception impacts the feeding preference of flies given a choice between 340mM sucrose and 340mM sucrose supplemented with cocaine (1.5mM or 3mM) or a known bitter compound, L-canavanine (10mM) (Lee et al., 2012). We observed a significant decrease in avoidance of cocaine and L-canavanine in Gr66a>Kir flies when compared to controls (UAS-Kir) (Figure 3a). The results of this experiment suggest that flies sense cocaine as a bitter tastant.

**Figure 3.**
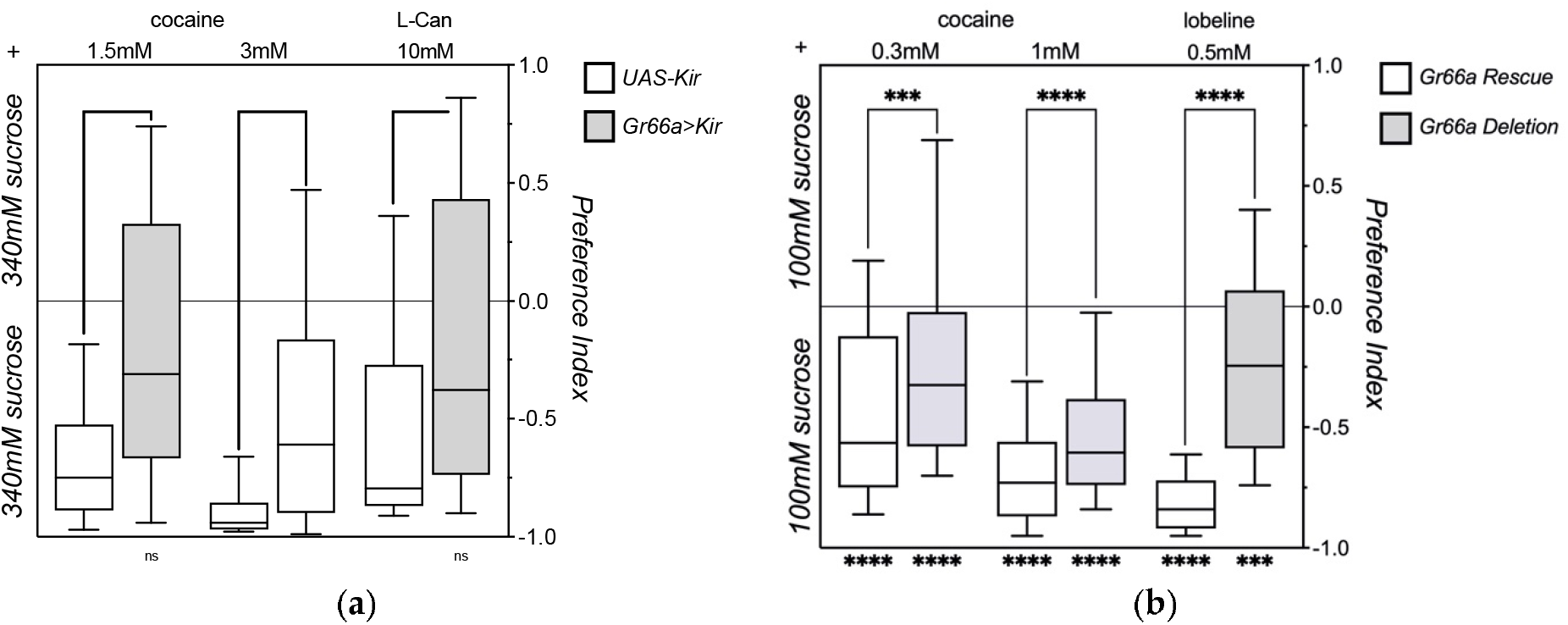
*Drosophila* Avoid Cocaine via G-Protein Coupled Receptors. Asterisks inside the graph’s frame indicate the P-value classification of two-tailed Mann-Whitney tests comparing median preference scores between genotypes. Asterisks beneath the frame of the graph show results of Wilcoxon signed-rank tests comparing the median preference to the hypothetical median of 0. **(a)** Silencing bitter taste neurons reduces cocaine avoidance in a two-choice fluorescent reading assay of preference. Control *UAS-Kir* flies (white bars) and bitter taste impaired *Gr66a>Kir* flies (grey bars) were wet starved for 6 hours, then allowed to choose between solutions containing 340mM sucrose or 340mM sucrose supplemented with cocaine. Compared *to UAS-Kir* control flies (white bars), bitter taste impaired flies (grey bars) display a significant decrease (two-tailed Mann-Whitney test) in avoidance for sucrose supplemented with 1.5mM cocaine (\*\*\*\**P* < 0.0001, n = 44, 44 groups of 3 flies), 3mM cocaine (\*\*\*\**P* < 0.0001, n = 44, 41), and the bitter compound L-canavanine (L-can) (\*\*\**P* < 0.0006, n= 61, 59; two-tailed Mann-Whitney U test). *UAS-Kir* control display significant avoidance for sucrose solutions supplemented with 1.5mM (\*\*\*\**P* < 0.0001, n = 44, Wilcoxon signed-rank test) and 3mM (\*\*\*\**P* < 0.0001, n = 41) cocaine, and 10mM of the bitter control compound L-canavanine (L-can) (\*\*\*\**P* < 0.0001, n = 62). In contrast, the *Gr66a>Kir* flies displayed reduced bitter avoidance, with significant aversion for 3mM cocaine (\*\*\*\**P* < 0.0001, n = 44), but not 1.5mM cocaine (*P* = 0.0998, n = 44) or 10mM L-canavanine (*P* = 0.0870, n = 59). **(b)** Deletion of the gustatory receptor subunit 66a gene (*Gr66a*) reduces cocaine avoidance in a two-choice fluorescent reading assay of preference. Gr66a rescue (white bars) and Gr66a deletion (grey bars) were wet starved for 18 hours, then allowed to choose between fluorescently labeled color counter-balanced solutions containing 100mM sucrose or 100mM sucrose supplemented with cocaine. Compared to *Gr66a Rescue* flies (white bars), *Gr66a Deletion* bitter taste mutants (grey bars) display a significant decrease (two-tailed Mann-Whitney test) in avoidance for sucrose supplemented with 0.3mM cocaine (***P < 0.0001, n = 128, 128 groups of 3), 1mM cocaine (****P < 0.0001, n = 144, 144 groups of 3), and the bitter compound lobeline (***P < 0.0001 n = 48, 48 groups of 3). Median preference scores for *Gr66a Rescue* flies were significantly less than a hypothetical median score of 0 (Wilcoxon signed-rank test) for sucrose solutions supplemented with 0.3mM cocaine (****P < 0.0001, n = 128 groups of 3), 1mM cocaine (****P < 0.0001, n = 144 groups of 3), and lobeline (****P < 0.0001, n = 48 groups of 3). Preference scores for *Gr66a Deletion* bitter taste mutants were also significantly less than 0 (Wilcoxon signed-rank test) for flies offered 0.3mM cocaine (****P < 0.0001, n = 128 groups of 3), 1mM cocaine (****P < 0.0001, n = 144 groups of 3)., and lobeline (***P = 0.0006, n = 48 groups of 3).

In *Drosophila*, gustatory neurons that mediate avoidant behavioral responses express ionotropic (Ir) clade receptors and G-protein like Gr receptors (Koh et al., 2015). Since silencing the bitter gustatory receptor neurons expressing Gr66a reduces cocaine avoidance, we asked whether the actual G-coupled protein receptor Gr66a was involved in *Drosophila* cocaine detection. To test this hypothesis, we used two fly lines with transgenes that rescue different portions of a 3.3kB deletion fragment that includes Gr66a and the two genes that flank Gr66a (Moon et al., 2006). One of these lines includes a transgene that rescues the two genes flanking Gr66a (Gr66a deletion), while the other includes a transgene that rescues all three genes from the 3.3kB fragment, including Gr66a (Gr66a rescue). We measured preference in another 30-minute FRAP, but increased the deprivation length to 18 hours to ensure that all flies in the assay would eat. Data from our previous experiments indicate that 18-hour starved flies offered solutions containing 340mM sucrose will gorge on the first solution they contact. We, therefore, lowered the concentration of sucrose offerings in the FRAP to 100mM to increase the probability that flies would sample each food source. Flies were given a choice of 100mM sucrose and 100mM sucrose supplemented with different concentrations of cocaine (0.3mM, 1mM) or lobeline (0.5mM), a bitter phytotoxin detected by Gr66a (Kim et al., 2017). We observed a significant reduction in avoidance of cocaine (0.3mM and 1mM) and lobeline (0.5mM) in Gr66a deletion flies compared to Gr66a rescue flies (Figure 3b). These results suggest that flies sense cocaine as an aversive tastant using G-coupled protein receptor complexes that include Gr66a.

### Cocaine Aversion Involves Peripheral Detection by the Legs of *Drosophila*

Because flies integrate signals from multiple taste organs to decide whether to initiate feeding behavior, we wondered if cocaine avoidance might occur prior to ingestion. In addition to the taste hairs found in the mouthparts of *Drosophila*, chemosensory neurons are also distributed along the wings, abdomen, and on the distal tarsal segments of the legs. Because taste hairs on *Drosophila* legs allow flies to interact with and sample a food source prior to consumption with the proboscis, we wondered if Gr66a-expressing neurons might provide *Drosophila* a peripheral mechanism for cocaine detection and avoidance. To answer this question, we performed a 2-choice assay of preference using the fly liquid-food interaction counter (FLIC).

The FLIC is a feeding assay that records the type and length of interactions that occur over time (Ro et al., 2014) since it allows discrimination between leg events (LE), which represent tasting, and proboscis events (PE), indicative of consumption (Figure 4a,b). To verify that the FLIC is an effective assay for measuring cocaine preference in *Drosophila*, we first performed a feeding assay with wild-type control flies. We starved flies for 18 hours, then transferred flies to the FLIC and offered them a choice between 100mM sucrose and 100mM sucrose supplemented with cocaine (0.02mM, 0.05mM, or 0.1mM). We selected lower concentrations of cocaine because control flies still showed strong aversion for 0.3mM cocaine in the FRAP (Figure 3b) and because the FLIC provides greater resolution than our blue dye feeding assay and the FRAP. It also allowed us to further characterize the dynamic range of *Drosophila* sensitivity to the effects of cocaine on feeding behavior.

**Figure 4.**
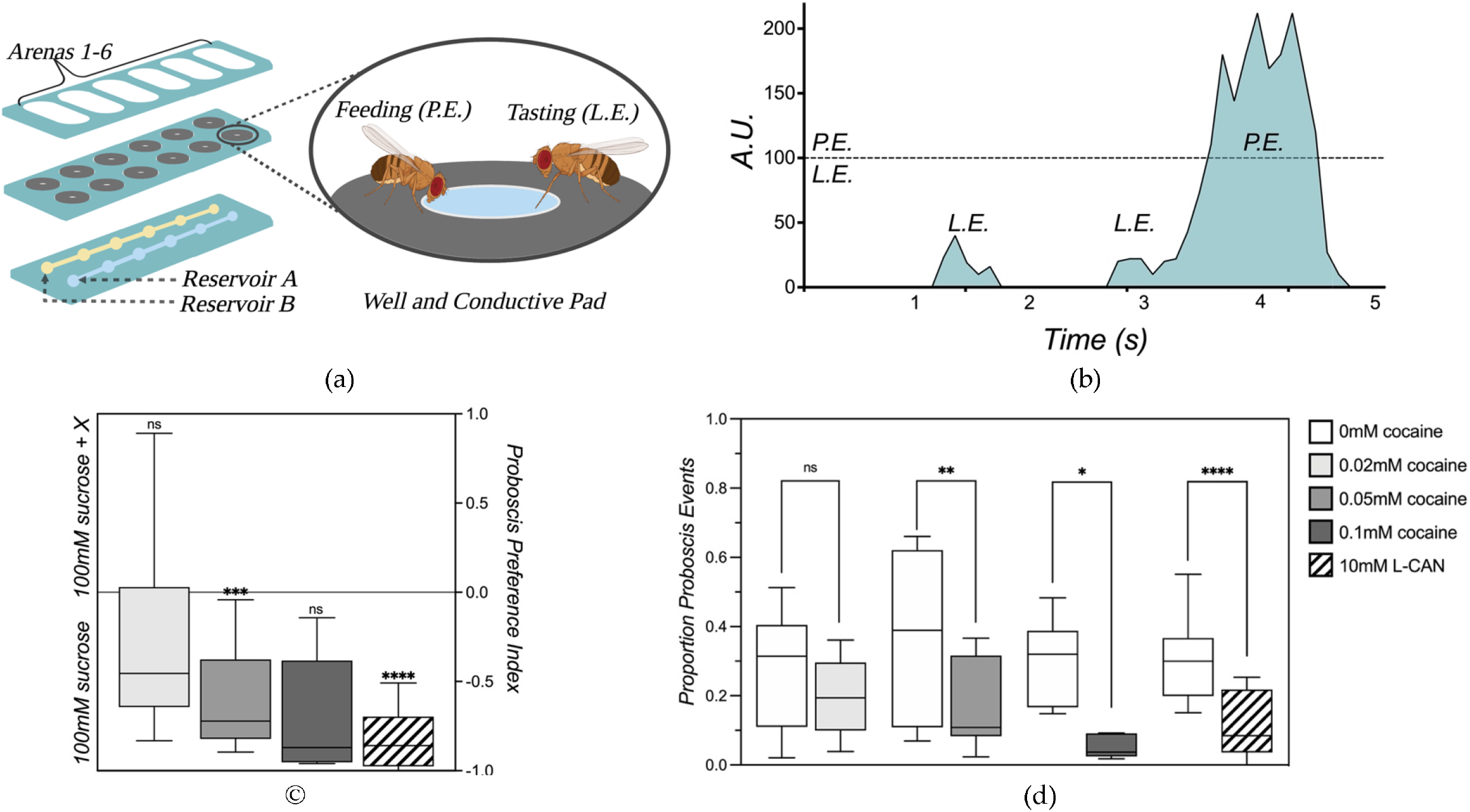
Cocaine is Aversive through a Peripheral Leg Mechanism. **(a)** A schematic of the 12 feeding wells from a single printed circuit board (PCB) used in the FLIC (1). Each FLIC board is fed by two reservoirs (A and B), where each reservoir runs the length of the board and supplies feeding solution to one lane of six wells. The PCB cover partitions the flic board into six arenas, each containing one well fed by reservoir A and one by reservoir B. When a fly standing on one of the conductive metal signal pads touches the feeding solution, they complete a voltage divider circuit (2). **(b)** Representative traces of FLIC data from a single well showing length and type of feeding events recorded over time. Leg events (L.E.) and proboscis events (P.E.) produce distinct changes in voltage that allow event discrimination based on the intensity of the normalized signal (A.U.). Values less than 100 represent leg events, and values over 100 represent proboscis events. Recordings from our experiments were used to generate readouts of activity in increments of 200ms, a resolution that allows identification of both single and clustered feeding events. **(c)** Flies avoid cocaine in a two-choice assay of preference in the fly liquid interaction counter (FLIC). After 18 hours of wet starvation, ten flies were loaded into each arena and allowed to choose between a solution of 100mM sucrose or 100mM sucrose supplemented with cocaine in a 30-minute assay of feeding. A proboscis preference index was calculated for each arena based on the difference in cumulative bout length for proboscis events at each well divided by the total length of proboscis events. A Wilcoxon signed-rank test was used to compare the median preference from each treatment to a hypothetical median of 0 (no preference). Cocaine produced a dose-dependent decrease in median preference, which is significantly deviates from 0 for sucrose solutions supplemented with 0.05mM (**P < 0.0002, n = 17 groups of 10), and 10mM L-canavanine (****P < 0.0001, n = 20, groups of 10). In contrast, median preference for flies offered 0.02mM cocaine did not significantly deviate from 0 (P = 0.1089, n = 17, groups of 10). Due to the strong aversion observed at 0.1mM cocaine, we only performed this experiment once, and, due to the low n, the proboscis preference was not significant for 0.1mM cocaine (P = 0.0312, n = 6, groups of 10). **(d)** Cocaine inhibits feeding initiation in the FLIC. Since all feeding events (P.E.) in the FLIC are preceded by tasting events (L.E.), the proportion of leg events that turn into proboscis events reflect food acceptance. We calculate this value for each well, determining the ratio of proboscis events to total events. A two-tailed Mann-Whitney test was performed to compare the median proboscis event proportion between pairs of wells from each treatment condition. A significant decrease in median proboscis event proportion was observed on wells supplemented with 0.05mM cocaine (*P < 0.0241, n = 14, 14 groups of 10, two-tailed Mann-Whitney), 0.1mM cocaine (**P < 0.0022, n = 6, 6 groups of 10 flies, two-tailed Mann-Whitney), and 10mM L-canavanine (***P < 0.0001, n = 6 groups of 5 flies, two-tailed Mann-Whitney). In contrast, we did not observe a significant difference in proboscis event proportion between wells when sucrose was supplemented with 0.02mM cocaine. Illustrations created with BioRender.com.

The cumulative duration of proboscis events, proportional to the volume ingested (Ro et al., 2014), recorded during the 30-minute FLIC assay was used to determine preference. We observed a dose-dependent decrease in proboscis preference for sucrose with cocaine, with significant avoidance for sucrose containing 0.05mM cocaine and 0.1mM cocaine (Figure 4c). The median proboscis preference index score for sucrose containing 0.02mM cocaine was not significantly different from 0 (Figure 4c). These results suggest that 0.02mM cocaine might be at the lower end of the dynamic range of *Drosophila* sensitivity to the bitter taste of cocaine in acute assays of feeding preference. Overall, the results of our time-based proboscis interaction preference measurements in the FLIC are consistent with our consumption-based measurements in the FRAP, suggesting that flies exhibit naïve cocaine avoidance, which increases dose-dependently.

Both the FRAP and the FLIC are useful for assaying preference. The FLIC, however, has the additional ability to measure feeding initiation for each interaction event with the food, enabling comparison of food palatability in a 2-well arena. In flies, the feeding behavior unfolds in a characterized sequence of events, with tasting events (leg interactions) preceding feeding events (proboscis interactions; see Figure 4b). In the FLIC, feeding initiation is reflected by the ratio of proboscis events to total events; i.e., how many interaction events with the food are ‘palatable enough’ to result in a proboscis/feeding event. Feeding initiation on the well with 100mM sucrose + 10mM L-canavanine, a bitter control substance, was much reduced compared to the other choice well of 100mM sucrose (Figure 4d). The same was true for 0.1mM and 0.05mM cocaine, while initiation for 0.02mM cocaine was not significantly different (Figure 4d). The decrease in the proportion of proboscis events observed for wells containing 0.05mM and 1mM cocaine suggests that cocaine is aversive via a mechanism of cocaine detection involving the legs, thus lowering the frequency of transition from leg into proboscis events.

To determine whether bitter gustatory receptors on the legs of *Drosophila* facilitate peripheral detection and avoidance of cocaine, we performed another FLIC experiment using Gr66a Deletion and Gr66a Rescue flies. We starved flies for 18 hours, then offered them a choice between 100mM sucrose and 100mM sucrose with cocaine (0.05mM or 0.1mM). We observed a significant reduction in avoidance of cocaine (0.05mM and 0.1mM) and L-canavanine (10mM) containing sucrose solution for Gr66a deletion flies compared to Gr66a rescue flies (Figure 5a). Next, we measured feeding initiation through the proportion of proboscis events to determine whether the reduction in cocaine avoidance observed in Gr66a deletion flies impacted cocaine perception by the legs. Gr66a rescue flies displayed a significant decrease in the proportion of proboscis events on wells containing 0.1mM cocaine and 10mM L-canavanine. In contrast, Gr66a deletion flies did not display a significant difference in the proportion of proboscis events between wells containing sucrose and wells containing sucrose with 0.1mM cocaine or 10mM L-canavanine. Surprisingly, Gr66a deletion flies given a choice between 100mM sucrose and 100mM sucrose with 0.05mM cocaine displayed a significant increase in the proportion of proboscis events, whereas no significant difference was observed in Gr66a rescue flies (Figure 5b). Together, these results suggest that cocaine inhibits feeding initiation in *Drosophila* by activating bitter gustatory receptors in the legs.

**Figure 5.**
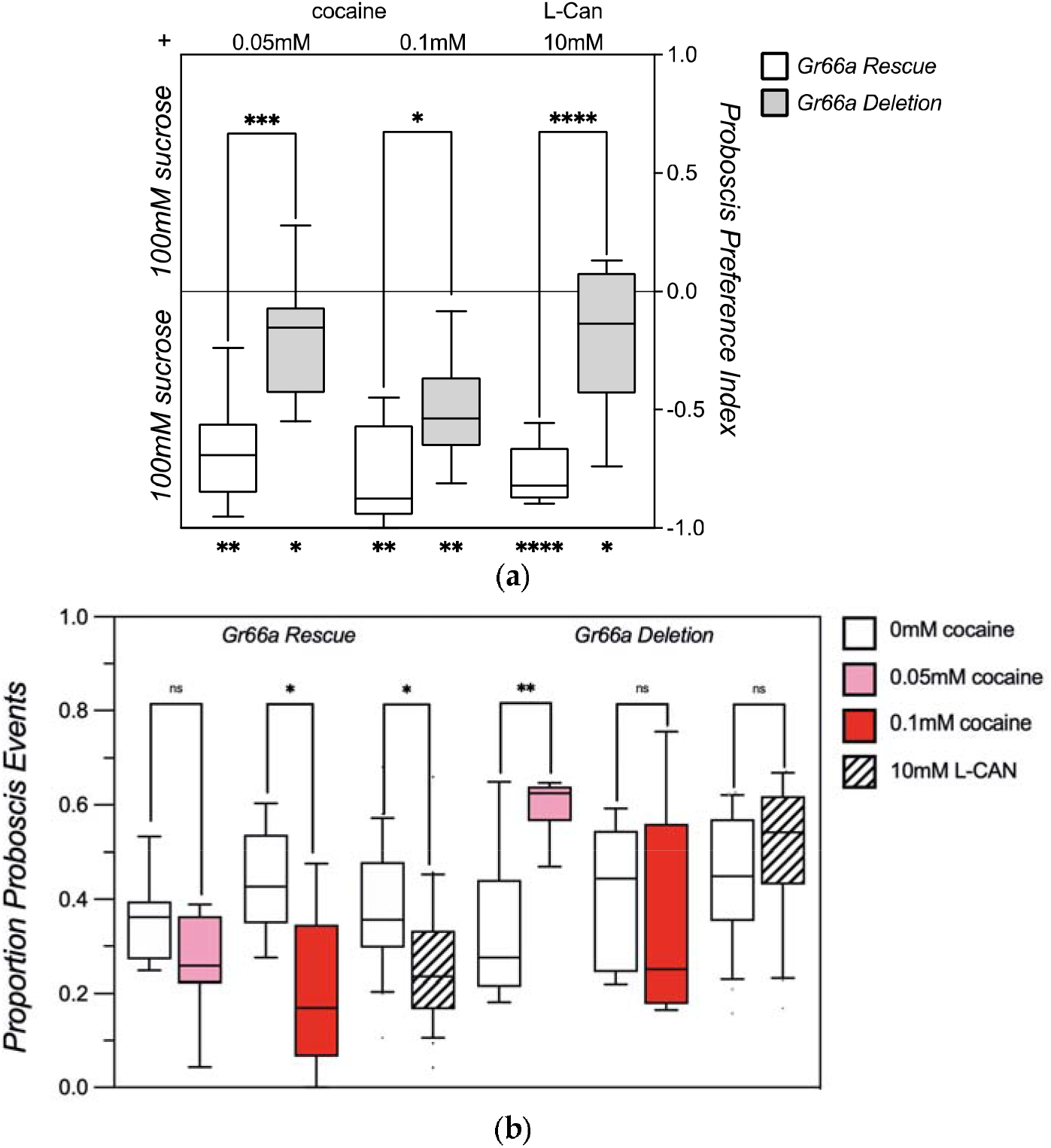
G-Protein Coupled Receptors are Essential for Leg-Mediated Detection and Avoidance of Cocaine. **(a)** *Gr66a Deletion* flies show decreased cocaine avoidance in the FLIC. All flies were wet starved for 18 hours, then transferred to FLIC boards by mouth-pipetting ten flies into each arena. Flies were allowed to choose between a solution of 100mM sucrose or 100mM sucrose supplemented with cocaine in a 30-minute assay of feeding. Asterisks inside the frame of the graph indicate the P-value classification for two-tailed Mann-Whitney tests comparing median preference scores between genotypes. Asterisks beneath the graph’s frame represent P-value classification for Wilcoxon signed-rank tests comparing the median proboscis preference index to a hypothetical median of 0. The median proboscis preference index score for Gr66a rescue flies was significantly less than 0 at −0.69 for 0.05mM cocaine (**P = 0.0039, n = 9 groups of 10 flies), −0.88 for 0.1mM cocaine (**P = 0.0039, n = 9 groups of 10 flies), and −0.82 for 10mM L-canavanine (****P < 0.0001, n = 21 groups of 10 flies). Median proboscis preference scores also deviated from 0 for Gr66a Deletion flies with a value of −0.15 for FLIC boards with 0.05mM cocaine (*P = 0.00391, n = 9 groups of 10 flies), −0.54 for 0.1mM cocaine (**P = 0.0078, n = 8 groups of 10 flies), and −0.1359 10mM L-canavanine (*P = 0.0158, n = 21 groups of 10 flies). The median proboscis preference index scores for *Gr66a Rescue* and *Gr66a Deletion* flies were compared for each treatment condition using the two-tailed Mann-Whitney test. Compared to *Gr66a Rescue* flies (white bars), *Gr66a Deletion* flies (grey bars) display a significant decrease in avoidance for sucrose supplemented with 0.05mM cocaine (***P < 0.0008, n = 9, 9 groups of 10), 0.1mM cocaine (*P < 0.0360, n = 8, 9 groups of 10), and the bitter compound L-canavanine (****P < 0.0001 n = 21, 21 groups of 10). **(b)** *Gr66a Deletion* increases initiation on sucrose supplemented with cocaine. A two-tailed Mann-Whitney test was performed to compare the median proboscis event proportion of wells in arenas from each treatment. A significant decrease in median proboscis event proportion was observed for *Gr66a Rescue* flies on wells with 0.1mM cocaine (*P < 0.0106, n = 9 groups of 10 flies, two-tailed Mann-Whitney) and 10mM L-canavanine (**P < 0.0052, n = 21 groups of 10 flies, two-tailed Mann-Whitney), but no difference was observed in arenas where cocaine was supplemented at 0.05mM (P = 0.0770, n = 9). In contrast, for *Gr66a Deletion* flies, we did not detect a significant difference in proboscis event proportion between wells when sucrose was supplemented with 0.1mM cocaine (P = 0.0921, n = 8 groups of 10 flies, two-tailed Mann-Whitney) or 10mM L-canavanine (P = 0.0001, n = 21 groups of 10 flies). Moreover, in arenas with 0.05mM cocaine we observed a significant increase in proboscis event proportion (**P < 0.0056, n = 9 groups of 10 flies).

### Cocaine Activates Gr66a Expressing Neurons on *Drosophila* Legs

Because our data from the FLIC indicated that *Drosophila* detect cocaine with their legs, we tested whether we could detect a response in sensory neurons through functional imaging using the genetically encoded calcium indicator GCaMP6s (Chen et al., 2013). To verify that we could reliably detect fluorescence in the legs of *Drosophila*, we first used the Gr66a driver to express enhanced green fluorescent protein (EGFP) in neurons expressing Gr-66a (Goentoro et al., 2006). We observed EGFP expression (Figure 6) in two bilateral pairs of neurons in the fifth tarsal segment and two bilateral pairs in the fourth tarsal segment of foreleg samples from adult *Drosophila* males. Next, we used the same Gr66a-Gal4 to drive the calcium indicator GCaMP6s (Chen et al., 2013). We confirmed expression by imaging the forelegs of adult Drosophila males, observing GCaMP6s fluorescence in the same regions of the distal tarsal segments identified during imaging of flies expressing EGFP (Figure 6). To functionally verify the response of bitter sensing gustatory neurons, we imaged the forelegs of adult *Drosophila* males and monitored GCaMP6s fluorescence during treatment with either sucrose, cocaine, or the bitter control compound denatonium (Figure 7). As expected, no change in baseline fluorescence intensity was observed after sucrose application, while incubation with denatonium increased GCaMP6s fluorescence, reflecting an increase in neuronal activation. We also observed a dose-dependent increase in GCaMP6s signal intensity in legs treated with different doses of cocaine. The results of our experiment provide functional verification of peripherical cocaine detection by *Drosophila* bitter gustatory neurons.

**Figure 6.**
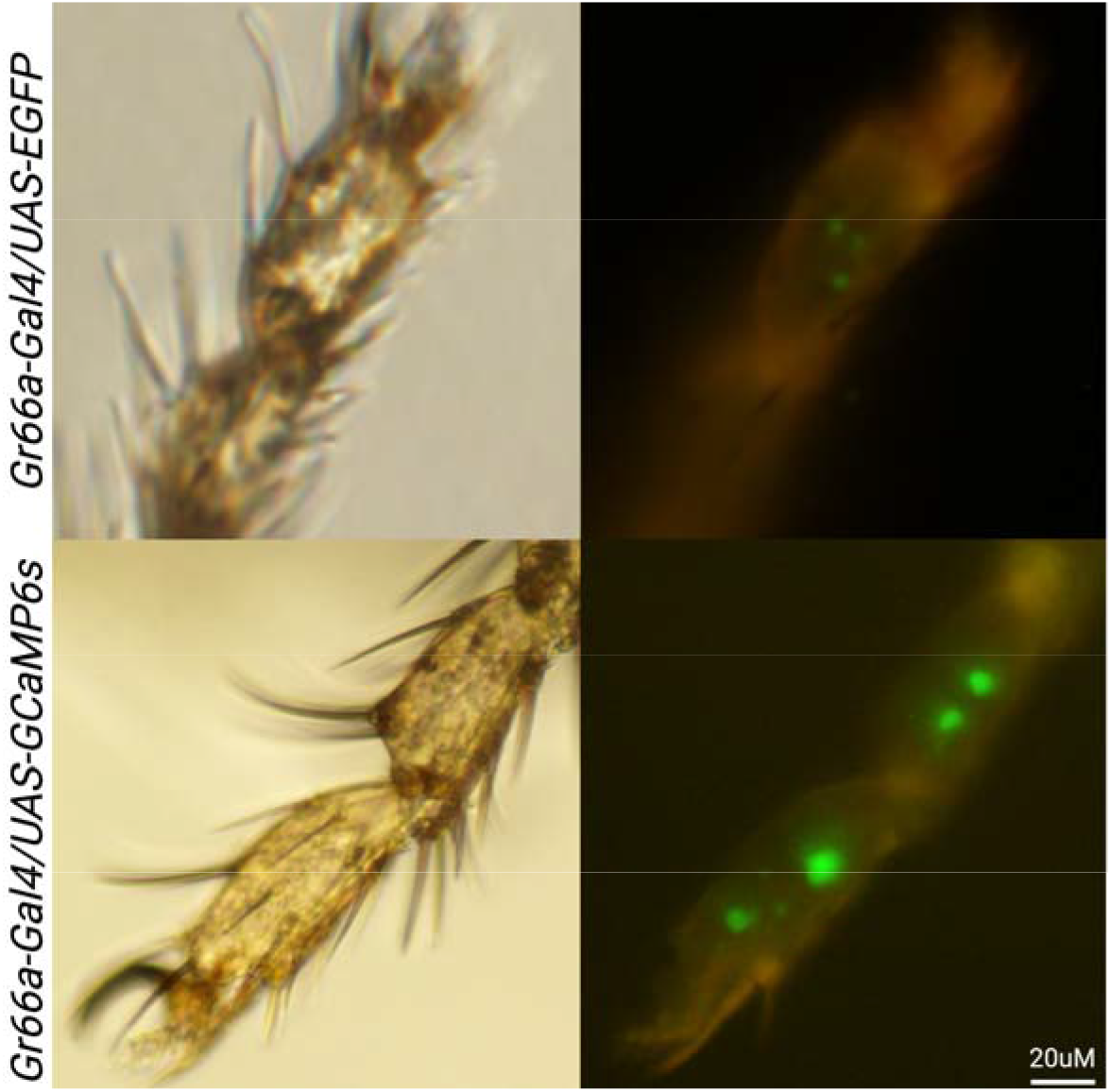
Gr66a-Gal4 is Expressed in the Distal Tarsal Segments of the *Drosophila* Foreleg. Representative images of fluorescence signal from *UAS-EGFP* (top right) and *UAS-GCaMP6* (bottom right) expressed in taste neurons of *Gr66a-Gal4* flies. Expression was observed in four bilateral pairs of bitter taste neurons, including two pairs in the fourth tarsal segment and two pairs in the fifth tarsal segment. All images were acquired using adult *Drosophila* males. Note, that we applied a higher gain and exposure for the *GCaMP6s* compared to the *EGFP* picture.

**Figure 7.**
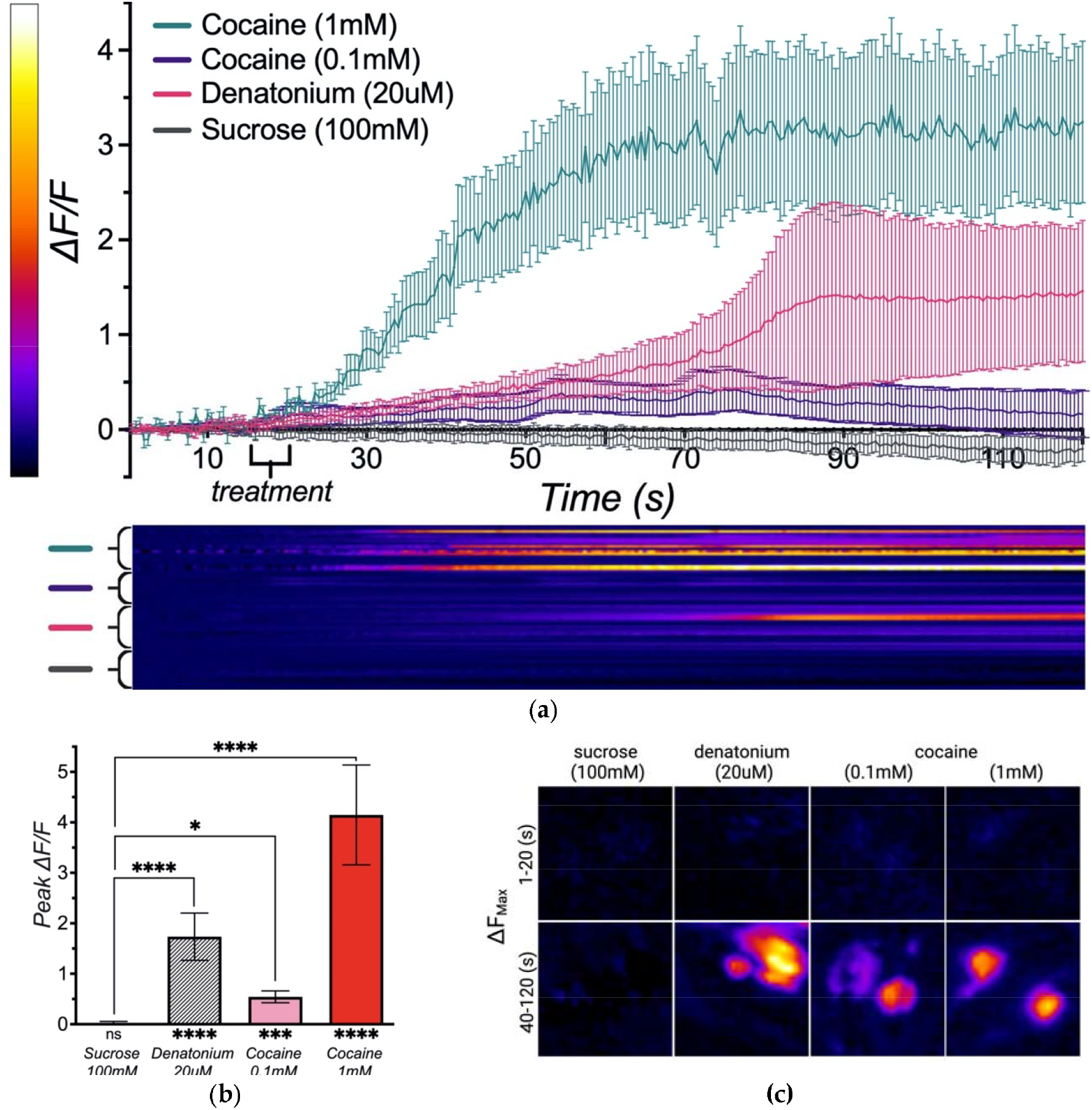
Cocaine is Detected by Taste Hairs on the Legs of *Drosophila*. **(a)** Increases in GCaMP6s fluorescence intensity were recorded in the forelegs of adult *Drosophila* males treated with 20uM denatonium (red trace with grey error bars), 1mM cocaine (red trace with teal error bars), and 0.1mM cocaine (red trace with light teal error bars). No increase in signal was recorded in forelegs treated with 100mM sucrose (black trace with grey error bars). The blue and red calibration bar to the left of the Y-axis represents the range of signal intensity observed during acquisition, and the heat map beneath the X-axis corresponds to the change in signal intensity over time in individual neurons from each treatment group. Plotted traces and error bars represent the mean change in signal intensity (ΔF/F) and the standard error of the mean (SEM) for neurons in each condition during the period 10 seconds before and 110 seconds after treatment. **(b)** Average peak intensity (ΔF/F) of neurons in each treatment condition. Kruskal–Wallis test with Dunnett’s post-hoc analysis was performed to compare the average peak intensity of sucrose treated control neurons (n = 17 neurons) to neurons treated with 20uM denatonium (****P = <0.0001, n = 18), 0.1mM cocaine (*P = 0.0201, n = 14), or 1mM cocaine (****P = <0.0001, n = 18). Asterisks beneath the graph’s frame represent P-value classification for Wilcoxon signed-rank tests comparing the median peak intensity (ΔF/F) to a hypothetical peak intensity of 0. The median ΔF/F for sucrose-treated controls was not significantly different than 0 (P = 0.7735). In contrast, median ΔF/F was significantly greater than zero for neurons treated with 20uM denatonium (****P = <0.0001), 0.1mM cocaine (***P = 0.0001), or 1mM cocaine (****P = <0.0001). **(c)** Max intensity projections of GCaMP6s fluorescence signal acquired before treatment (1-10s, top) and after treatment (20-120s, bottom).

## Discussion and Conclusions

Our experiments show that cocaine activates bitter-sensing peripheral gustatory neurons, enabling peripheral detection and pre-ingestive avoidance. Our results explain why developing a protocol for preferential cocaine self-administration in flies has proved difficult. In *Drosophila*, taste receptors distributed across internal and external tissues have unique roles in modulating feeding responses (Dunipace et al., 2001; Freeman and Dahanukar, 2015). Grs in the wings are important for locating food sources (Raad et al., 2016), whereas Grs in the tarsal segments are important to assess food before ingestion and impact food acceptance and preference (Meunier et al., 2003). Bitter Grs on the labellum influence food acceptance, preference, and consumption (Hiroi et al., 2004; Weiss et al., 2011; Harris et al., 2015), and projections from pharyngeal taste neurons to motor neurons enable Gr-mediated signaling to directly influence feeding bout length (Basiri and Stuber, 2016). Unique sensory information is encoded by activation of tarsal or labellar Grs, suggesting that tissue-specific disruption of Gr activation can significantly impact learning and behavior, including drug-taking behavior (Devineni et al., 2021)

*Drosophila* bitter Gr complexes vary in selectivity, and are either broadly or narrowly tuned to respond to aversive tastants (Weiss et al., 2011). However, although different bitter tastants can activate distinct receptors, *Drosophila* display minimal ability to discriminate between different bitter compounds (Masek and Scott, 2010). Furthermore, concentration-dependent shifts in neuronal and behavioral responses are similar across most bitter compounds, suggesting bitterness is perceived on a quantitative rather than qualitative scale (Hiroi et al., 2004; Sellier et al., 2011). This lack of discernment may be useful for developing *Drosophila* models of substance abuse, as reducing the sensitivity to bitterness might also be effective in reducing avoidance of plant toxins, such as cocaine.

We have demonstrated that flies naively avoid cocaine, which activates bitter-sensing neurons in the leg. The relationship between taste sensitivity and drug consumption observed in our experiments is also observed in humans. For example, bitter taste perception also modulates drug consumption in humans, and polymorphisms in bitter receptor proteins have been associated with the risk of developing alcoholism (Edenberg and Foroud, 2006; Hinrichs et al., 2006). Furthermore, genetic variation in bitter taste-sensing type-2 receptors (TAS2R) is associated with differences in average alcohol consumption (Duffy et al., 2004; Lanier et al., 2005), total alcohol consumption, and taste sensation and liking of other phytochemical-containing beverages (Hayes et al., 2011). Distinct taste receptor polymorphisms have also been associated with binge-like drinking behavior (Wang et al., 2007) and higher daily alcohol intake (Dotson et al., 2012). These studies highlight the relationship between drug use and taste receptor genes in humans, demonstrating that single nucleotide polymorphisms can influence the risk of substance use and dependence (Hinrichs et al., 2006).

While our data indicate that Gr66a is involved in cocaine detection in *Drosophila*, the additional Gr subunits involved in cocaine detection are still unknown. (Ling et al., 2014). Functional gustatory receptors found in *Drosophila* taste neurons are trimeric complexes composed of different combinations of Gr subunit proteins (Kim et al., 2017), whose variation can lead to cell-specific differences in chemical sensitivity (Kwon et al., 2014; Ling et al., 2014). Twenty-eight bitter-sensing gustatory receptor subunits are expressed on the legs of *Drosophila*, but only a subset is expressed in each bitter-sensing neuron (Kwon et al., 2014; Ling et al., 2014). Future experiments should attempt to characterize the remaining Gr subunits that facilitate cocaine detection and avoidance in *Drosophila*.

While characterizing the components of a peripheral cocaine receptor will help identify methods for reducing cocaine avoidance in *Drosophila*, it is still possible that peripheral cocaine detection by tarsal taste receptors is only one component of *Drosophila* reluctance to consume cocaine. Many bitter compounds are detected by multiple Gr complexes, and it is possible that other Grs in the labellum or pharynx might still act to reduce cocaine consumption (Kim et al., 2017). Specifically, Gr66a deletion significantly reduces, but does not eliminate, cocaine avoidance in *Drosophila*. Whether residual cocaine avoidance is due to taste or some other noxious component of cocaine ingestion remains to be determined. Determining how gustatory receptors in other taste organs contribute to cocaine detection and avoidance will help answer this question. A complete understanding of the gustatory components involved in cocaine detection will help elucidate the basis of cocaine avoidance in *Drosophila* and support the development of a fly model of cocaine use disorder.

## Materials and Methods

### *Drosophila* Stocks

Behavioral experiments were performed with male flies raised in 12:12 hour L:D conditions at 25°C with 70% humidity on standard cornmeal/molasses food. Flies used in behavioral experiments were 3–6 days of age at the start of the experiments. Male *w* Berlin* flies were chosen as a genetic wild-type strain for all preliminary experiments of consumption, preference, and survival. For assays requiring food deprivation, flies were kept in vials containing 0.7% water agar for 6 or 18 hours. Transgenic flies expressing the genetically encoded calcium indicator GCaMP6s in Gr66a-positive taste neurons were generated by crossing *Gr66a-Gal4* flies (w[*]; *P{w[+mC]=Gr66a-GAL4.D}2*, extracted from RRID:BDSC_28801) with GCaMP6s flies (*w[1118]; P{y[+t7.7] w[+mC]=20XUAS-IVS-GCaMP6s}attP40*, RRID:BDSC_42746). Fly lines for taste experiments were obtained from the Bloomington Stock Center and include *w[*]; P{w[+mC]=Gr66a[+t8]}2; Df(3L)ex83* RRID:BDSC_35528 rescue of Gr66a, *w[*]; P{w[+mC]=CG7066[t7]}2; Df(3L)ex83* RRID:BDSC_28804 non-rescue of Gr66a, and *w[*]; Df(3L)ex83* RRID:BDSC_25027 the 3.3kb Df of 3 genes, incl Gr66a

### Chemicals and Reagents

Cocaine-HCl was provided by the National Institute on Drug Abuse under Drug Enforcement Administration license # RR0499699 (Rothenfluh).

### Blue Dye Feeding Assay

Male flies, 3-6 days old, were collected under CO2 anesthesia and allowed to recover for 24 hours in vials with standard cornmeal food. After recovery, flies were wet starved for 6, 12, or 18 hours in 0.7% water agar vials. Following starvation, flies were transferred to feeding vials with strips of filter paper (7 × 1.75cm) coated in 350ul feeding solution. All feeding solutions contained 0.3% (v/v) blue food dye (FD&C Blue Dye no. 1) and sucrose, or sucrose with cocaine, lobeline, or L-canavanine. After a 4-minute assay of feeding, flies were transferred to empty vials and frozen at −80°C. Flies that ate were identified under a stereomicroscope based on the presence of blue dye and transferred to Eppendorf tubes in groups of five. 50ul of water was added to each tube, and flies were homogenized with a small battery-powered pestle. Homogenates were spun down for 1 minute at 14,000rpm with the lid hinge oriented outward, then for 4 minutes at 14,000rpm with the lid hinge oriented outward. Centrifugation results in a pellet of tissue, leaving a supernatant of blue-dyed solution under a thin lipid layer. A NanoDrop spectrophotometer was used to measure absorbance on 2ul samples of the dyed supernatant, with special care being taken to avoid the transfer of the topmost lipid layer. Absorbance was measured at 630nm and 700nm to quantify consumption. Feeding volume for each fly was calculated using the formula nL eaten = (OD 630nm - 1.1*OD 700 nm) *CF, where CF is a conversion factor specific to the 3% Blue#1 stock solution.

### Fluorometric Reading Assay of Preference

Naive cocaine preference was tested in a 30-min two-choice fluorometric plate-reading assay described previously (Peru y Colón de Portugal et al., 2014), with some modifications. Male flies were collected in groups of 35 under CO2 anesthesia and allowed to recover for 24 hours in vials with standard cornmeal food. After recovery, flies were transferred to starvation vials containing 0.7% water agar and food deprived for 6 or 18 hours. Before the experiment, we prepared 60-well plates offering a choice between two equivalent sucrose solutions (340mM or 100mM), where one solution was supplemented with cocaine, lobeline, or L-canavanine. The feeding solutions in each plate were labeled with distinct fluorescent dyes (0.005% rhodamine B and 0.003% fluorescein) to determine relative consumption. To control for color bias, we ran two plates for each pair of feeding solutions, where the solutions in each plate were reciprocally dyed. 100ul of 0.005% rhodamine B sucrose solution and 100 ul of 0.003% fluorescein sucrose solution were mixed to obtain a standard 50/50 dye solution, which was distributed in 20 wells of a 60-well plate (10 ul per well). For all other plates, 100ul of 0.005% rhodamine B liquid sucrose solution and 100 ul of 0.003% fluorescein liquid sucrose solution were added across 20 wells in a staggered, offset pattern, including wells C1-C10 and E1-E10. Special care was taken to avoid filling wells underneath the hole in the lid through which flies are loaded into the plate.

Following starvation, flies were gently transferred to 60-well plates using a mouth pipette or a modified 10ml starvation pipette. After loading the flies, the holes in the lid of each plate were covered with tape, the plates were turned upside down, and the flies were allowed to feed for 30 minutes. After the assay, the plates were transferred to the −80°C freezer to flash freeze flies. We then performed whole-fly analysis with 24 flies from each plate in a Fluoroskan Ascent plate reader. We used a 384-well Greiner plate, suspending three flies in each well with 60uL of water (8 wells per plate). Next, we loaded the plates into the Fluoroskan and measured the relative emission intensity of each dye by scanning with filter pairs at 485/527 and 542/591, which correspond to fluorescein and rhodamine B, respectively. These data were used to generate a ratiometric index of consumption representing a relative preference for each solution.

### Fly Liquid Interaction Counter Assay

Fly Liquid Food Interaction Counter (FLIC) assays were performed as previously described (Ro et al., 2014) with some modifications. Male flies, 3-6 days old, were collected with CO2 anesthesia at least 24 hours before each experiment. Flies were food-deprived for 18 hours in vials with 0.7% agar to prevent dehydration. After starvation, we used a mouth pipette to transfer ten flies to each arena in the FLIC for a 30-minute assay of preference. During this period, flies in each area had free access to two feeding wells, one containing a 100mM sucrose solution and one containing 100mM sucrose solution supplemented with either cocaine (0.02, 0.05, 0.1mM) or L-Canavanine (10mM). The number and duration of leg events (LE) and proboscis events (PE) for each well in the *Drosophila* feeding monitor were recorded using the FLIC monitor software. PE and LE were defined as having a peak signal amplitude greater than or less than 100AU, respectively. Raw data from each experiment were processed using custom-built software to analyze proboscis preference, event preference, and percent of proboscis events.

As proboscis interactions are indicative of feeding events, the cumulative length of proboscis events at each feeding well provides an indirect measure of feeding activity. The proboscis preference index represents the relative interaction time between two wells and provides an indirect measure of feeding preference. Proboscis preference was calculated as the difference in interaction time between two wells divided by the total interaction time. Results of this analysis give a value between◻+◻1 and◻−◻1, with +1 representing absolute preference, −1 representing total avoidance, and a value of 0 indicating no preference. Event preference was calculated using the same formula but is based on the count of events rather than the length of events. The proportion of proboscis events to total events was used to determine acceptance for each well.

### Calcium Imaging Gustatory Neuron Responses in *Drosophila* Legs

Calcium imaging taste response in *Drosophila* legs was performed as previously described (Miyamoto et al., 2013) with some modifications. Flies were anesthetized with CO_2,_ and forceps were used to remove the legs by cutting between the femur and tibia. Double-sided tape was used to fix the tibia to a slide, and another piece of tape was used to secure the leg, leaving the distal 4th and 5th tarsal segments exposed over the glass. The exposed tarsal segments were covered with water, and the slide was transferred to the microscope for imaging. The legs were imaged for 30-60 seconds to establish baseline fluorescence, then treated with either sucrose (100mM), denatonium (20uM), or cocaine (0.1mM, 1mM). Aliquots of each treatment compound were mixed at concentrations of 10x or 20x, and 100ul or 200ul were added to each prep during treatment. Imaging was performed with an Olympus CKX53 inverted culture scope using a 40x objective and an X-Cite 120q fluorescence illumination light source. The preparation was excited with a mercury arc lamp using the CKX3-B-excitation fluorescence mirror unit with an excitation filter (BP460-490C), dichroic mirror (LP500), and emission filter (520IF). Image acquisition was performed using an Olympus DP72 microscope digital camera with DP Manager software (CellSens). Acquisition settings for representative images were optimized to enhance structural aspects of tarsal segments and support visualization of Gr66a-positive neurons. For calcium imaging, images were acquired at 2fps with an exposure time of 500ms and at the minimum setting for light sensitivity (ISO 200).

### Image Processing and Analysis

Image sequences for each leg (240 frames) were processed using ImageJ. Images were cropped to include the fourth and fifth tarsal segments, and XY-drift was corrected with TurboReg based on an average intensity Z-projection. ROIs were identified by analyzing particles in a standard deviation Z-projection generated from the registered image after applying the triangle auto-threshold method. The ROI manager was used to add ellipses to regions adjacent to cells to cells of interest to measure background fluorescence. Intensity measurements for each frame were used to plot changes in intensity over time. Twenty frames (10s) were analyzed to establish a baseline ΔF/F. Solutions were applied between frames 20-30 (15-20s), and responses were analyzed across the last 200 frames (100s). The change in fluorescence intensity (ΔF) relative to baseline fluorescence intensity (F) was used to plot ΔF/F for each acquisition series. Untreated cells were imaged to measure photobleaching. Results were fit to an exponential decay function and applied to correct signal intensity across each treatment condition.

### Data Analysis and Statistics

Statistical analyses were performed using GraphPad Prism 9. Data from feeding experiments were not assumed to follow a normal distribution; therefore, preference and consumption experiments were analyzed using non-nonparametric statistics. The two-tailed Mann-Whitney U test was used to determine the significance of pairwise comparisons in the blue-dye feeding assay of consumption, the FRAP, and FLIC. The Wilcoxon signed rank test with a two-tailed P-value was used in the FRAP and FLIC to compare the median preference score of each group to a hypothetical median of zero. Box plots in all consumption and preference experiments represent median values and interquartile range, with whiskers depicting 10^th^ and 90^th^ percentiles.

## Acknowledgements

We thank the members of the Rodan and Rothenfluh labs for continued discussion and suggestions. This work was supported by the NIH: the National Institute of Diabetes and Digestive and Kidney Diseases (R01DK110358 to A.R.R.), the National Institute on Drug Abuse (Grants R21DA049635 and R21DA040439 to A.R.), and the National Institute on Alcohol Abuse and Alcoholism (Grant R01AA026818 and R01AA019536 to A.R.).

